# The Role of Conjunctive Representations in Regulating Actions

**DOI:** 10.1101/2020.04.30.070227

**Authors:** Atsushi Kikumoto, Ulrich Mayr

## Abstract

Action selection appears to rely on conjunctive representations that nonlinearly integrate task-relevant features (Kikumoto & Mayr, 2020). We test here the corollary hypothesis that such representations are also intricately involved during attempts to stop an action—a key aspect of action regulation. We tracked both conjunctive representations and those of constituent rule, stimulus, or response features through trial-by-trial representational similarity analysis of the EEG signal in a combined, rule-selection and stop-signal paradigm. Across two experiments with student participants (*N* = 57), we found (a) that the strength of decoded conjunctive representations prior to the stop signal uniquely predicted trial-by-trial stopping success (Exp. 1) and (b) that these representations were selectively suppressed following the onset of the stop signal (Exp. 1 and 2). We conclude that conjunctive representations are key to successful action execution and therefore need to be suppressed when an intended action is no longer appropriate.

**Statement of Relevance:** Some theorists have posited that as a necessary step during action selection, action-relevant features need to be combined within a conjunctive representation that is more than the sum if its basic features. Consequently, such representations should also play a critical role when trying to stop an intended action—a key aspect of self-regulation. However direct evidence of conjunctive representations has been elusive. Using a method for tracking both conjunctive and basic-feature representations on a trial-by-trial basis in the EEG signal, we show that the stronger the conjunctive representations, the harder it was to stop the intended action. Furthermore, the stopping process also selectively reduced the strength of conjunctive representations. These results further our knowledge about action regulation by showing that conjunctive representations are a necessary precursor for carrying out actions successfully and for that reason also need to be the target of self-regulatory stopping attempts.

---

Even simple goal-directed actions, such as kicking a soccer ball to a teammate, rely on various sensorimotor features—the location of the ball, the presence of opponent players and teammates, as well as on abstract rules (e.g., “kick softly when the grass is wet”). In traditional, stage-type information processing models, such different task-relevant features are handled independently, and in a serial, feed-forward manner Donders (1969); (Kornblum, Hasbroucq, & Osman, 1990; Posner & Mitchell, 1967; Sanders & Sanders, 2013; Sternberg, 1969). Alternatively, there are models in which all relevant features are combined within a common representational space during selection. Specifically, event-file theory (Hommel, 1998, 2019; Hommel, Müsseler, Aschersleben, & Prinz, 2001; Schumacher & Hazeltine, 2016) posits that an action becomes executable only once all task-relevant features are integrated within conjunctive representations, also referred to as event files. Moreover, recent research in nonhuman primates indicates that neurons with nonlinear, mixed selectivity response properties integrate various aspects in a conjunctive manner and play a critical role in flexible control of task-relevant information (Parthasarathy et al., 2017; Rigotti et al., 2013; Stokes et al., 2013).

If conjunctive representations are necessary, and maybe even sufficient precursors of goal-directed behavior, it follows that the pathway towards regulating a given action also needs to lead through these representations. For example, when in the above soccer scenario, an opponent defender suddenly blocks the goal, the intended kicking action has to be quickly canceled. The cognitive and neural underpinnings of such response inhibition have been well characterized using variants of the stop-signal paradigm (Aron, Robbins, & Poldrack, 2014; Logan & Cowan, 1984; Swann et al., 2009; Verbruggen et al., 2019; Wessel, 2019). Yet, it is currently an open question how stopping affects the different representations that underlie planned actions (e.g., of stimuli, responses, or rules). For example, in theory, the stopping process might occur by suppressing solely response representations that directly link to motor control pathways (Coxon, Stinear, & Byblow, 2006; Duque, Greenhouse, Labruna, & Ivry, 2017; Greenhouse, Sias, Labruna, & Ivry, 2015; Labruna et al., 2014), while leaving other task-relevant representations intact. However, assuming conjunctions are indeed critical for action control, the cancellation of an initiated action should require the suppression of the entire, integrated representation.

To test this hypothesis, it is necessary to track the various relevant representations of task-relevant features concurrently while actions are selected and regulated. Recently, Kikumoto and Mayr (2020) applied time-resolved representational similarity analysis (RSA, Kriegeskorte, Mur, & Bandettini, 2008) to the EEG signal in order to track both conjunctive and constituent feature representations during rule-based action selection in humans. These analyses indeed revealed conjunctive representations that integrated action rules to specific sensory/motor settings throughout the entire selection period. Moreover, the strength of conjunctions was a robust and unique predictor of trial-to-trial variability in RTs—as one would expect if conjunctive representations are necessary and sufficient conditions for action execution.

To directly test the hypothesis that the pathway to canceling an action leads through the corresponding conjunctive representation, we combined here a rule-based action selection task (Fig. 1ab, Kikumoto & Mayr, 2020; Mayr & Bryck, 2005) with an occasional stop signal. In Exp. 1, the stop-signal timing was adjusted via an adaptive tracking procedure, based on participants’ trial-to-trial stopping accuracy to achieve around 50% stopping success. Our main goal here was to test the prediction that the strength of conjunctions prior to stopping, inversely predicts stopping success and we also aimed to provide initial information about which representations are targeted by the stopping process. In Exp. 2, the stop signal was presented 100 ms after the stimulus onset, which is early enough for successfully stopping actions in most trials. Here, our main goal was to clearly characterize the consequences of successful stopping of actions on task-relevant representations and specifically test the prediction that conjunctive representations are selectively suppressed following the stop signal.

**Fig. 1.**
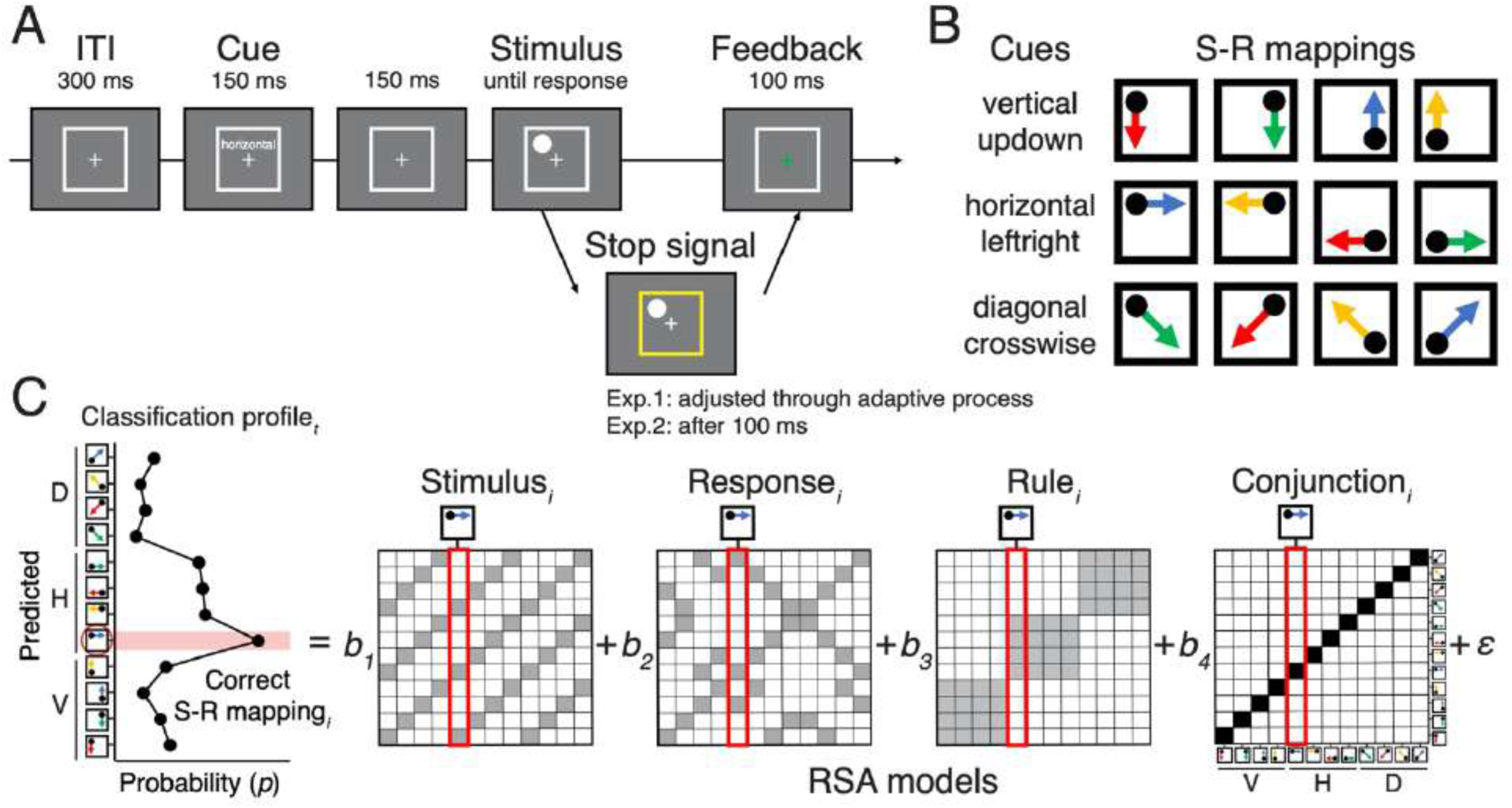
Task design and analytic procedure. (A) Sequence of trial events in the combined rule-selection/stop-signal task for both Exp. 1 and 2. (B) Spatial translation rules mapping specific stimuli to responses. Two different cue words were used for each rule to disambiguate between cue and rule-level representations. (C) Schematic steps of the representational similarity analysis. The raw EEG signal was decomposed into activity in specific frequency-bands via time-frequency analysis (see *EEG recordings and preprocessing* and *Time-Frequency Analysis*). For each sample time (*t*), a scalp-distributed pattern of EEG power was used to decode the specific rule/stimulus/response configuration of a given trial, producing a set of classification probabilities for each of the possible configurations. The profile of classification probabilities reflects the similarity structure of the underlying representations, where similar action constellations are more likely to be confused. The figure shows as an example classification probabilities for a case where both a unique conjunction and rule information are expressed (peak at the correct S-R mapping, plus confusion to other instances with the same rule). For each trial and timepoint, the profile of classification probabilities was regressed onto model vectors as predictors that reflect the different, possible representations. In each matrix of model vectors, the x-axis corresponds to the correct constellation for the decoder to pick, and the y-axis shows all possible constellation. The shading of squares indicates the theoretically predicted classification probabilities (darker shading means higher probabilities). The coefficients associated with each predictor (i.e., *t*-values) reflect the unique variance explained by each of the constituent features and their conjunction.

## Materials and Methods

### Participants

A convenience sample of 64 students of the University of Oregon participated after signing an informed consent following a protocol approved by the University of Oregon’s Human Subjects Committee in exchange for the compensation of $10 per hour and additional performance-based incentives. Participants with excessive amount of EEG artifacts (i.e., more than 35% of trials; see *EEG recordings and preprocessing* for detail) were removed from further analysis. As a result, we retained 36 out of 38 participants for Exp. 1, and 24 out of 26 participants for Exp. 2. In Exp. 1, three participants were further excluded because of failures to stop in excess of 75%. Samples sizes for the two experiments were based on our previous work (Kikumoto & Mayr, 2020), in which we obtained very robust results with sample sizes around 20 participants. For Exp. 1, we increased the target sample size to 36 participants, because here we were interested in a differentiation between failed and successful stop trials.

### Stimuli, Tasks and Procedure

Participants were randomly cued on a trial-by-trial basis to execute one of the three possible actions rules (Fig. 1a, Mayr & Bryck, 2005). Based on the cued rule, participants responded to the location of a circle (1.32° in radius) that randomly appeared in the corner of a white frame (6.6° in one side) by selecting one of the four response keys that were arranged in 2 x 2 matrix (4, 5, 1, and 2 on the number pad). For example, the vertical rule mapped the left-top dot to the bottom-left response as a correct response. Two different cue words (e.g., “vertical” or “updown”) were used for each rule (i.e., 66.6 % switch rate).

In 33.3% of trials, the stop signal (i.e., a yellow frame; Fig. 1a) indicated to participants that the planned action had to be cancelled. Stop-trials were counted as successful when participants did not make any responses within 800 ms time-window following the stop-signal onset. In Exp. 1, the interval between the stimulus and stop-signal onset (i.e., the stop-signal delay or SSD) was adjusted using an adaptive tracking method based on participants’ trial-to-trial stopping success. Specifically, individuals’ SSDs varied between 0 ms to 800 ms counting from the onset of the stimulus and starting with 100 ms at the beginning of session. Correct/incorrect stop trials increased/decreased SSDs by the step size that was randomly selected from 11.8 ms, 23.5 ms, or 35.3 ms for each trial. In Exp. 2, the stop signal appeared 100 ms after the stimulus onset. Go trials lasted until either the response was executed; stop trials lasted either until the 800 ms response window expired, or until a response was recorded.

There were two practice blocks and 200 and 250 experimental blocks for Exp. 1 and 2 respectively. Each block lasted 15 seconds, within which participants were instructed to complete as many trials as possible. Trials that were initiated within the 15 second block duration were allowed to complete. The average number of go-trials and stop-trials were 1576 (SD = 162) and 773 (SD = 75) for Exp. 1, and 1378 (SD = 91) and 685 (SD = 33) for Exp. 2. Throughout the experimental session, participants were reminded to respond as accurately and fast as possible and refrain from waiting for the stop signal. In Exp. 1, participants were instructed that the adaptive tracking procedure would make it easier to stop on some trials and more challenging on others. Participants were given a performance-based incentive for trials with RTs on go-trials faster than the 75th percentile of correct responses in the preceding blocks when 1) the overall accuracy in go-trials was above 90 percent and 2) there were more than 5 completed trials in a given block. While performing the task, participants were asked to rest the index finger in the center of the four response keys at the start of each trial (i.e., no lateralization of response sides). At the end of each trial, feedback (a green fixation cross for correct and a red cross for correct trials) was presented based on the accuracy of responses in go-trials or on correct stopping in stop-trial. At the end of each block, the number of completed trials, the number of correct responses in go/stop-trials, and the amount of earned incentives based on the speed of responses in go-trials, were presented as a feedback. All stimuli were created in Matlab (Mathworks) using the Psychophysics Toolbox (Brainard, 1997) and were presented on a 17-inch CRT monitor (refresh rate: 60 *Hz*) at a viewing distance of 100 cm.

### Stop-signal Reaction Time

In Exp. 1, we computed individuals’ stop-signal reaction time (SSRT), according to the integration method as specified by Verbruggen et al. (2019). First, for each quantile bin of SSDs (Fig. 3b), the mean SSDs and the proportion of successful stop trials (*p*(respond|signal)) were calculated. Then, the matching go RTs were defined in each SSD bin by taking the *n*th RT in the rank ordered go-trial RTs (including all go-trials), where *n* is defined by multiplying the number of RTs in the distribution by the probability of responding, *p*(respond|signal) or unsuccessful stopping, for each SSD bin. Within each SSD bin, SSRT was calculated by subtracting the corresponding SSD from the matching go RT, then scores from 6 SSD bins were averaged within individuals to obtain a single metric of SSRT for each individual. For all participants, failed-stop RTs were faster than correct go RTs.

### EEG recordings and preprocessing

Scalp EEG activities were recorded from 20 tin electrodes on an elastic cap (Electro-Caps) using the International 10/20 system. The 10/20 sites F3, Fz, F4, T3, C3, CZ, C4, T4, P3, PZ, P4, T5, T6, O1, and O2 were used along with five nonstandard sites: OL halfway between T5 and O1; OR halfway between T6 and O2; PO3 halfway between P3 and OL; PO4 halfway between P4 and OR; and POz halfway between PO3 and PO4. Electrodes placed ~1cm to the left and right of the external canthi of each eye recorded horizontal electrooculogram (EOG) to measure horizontal saccades. To detect blinks, vertical EOG was recorded from an electrode placed beneath the left eye and reference to the left mastoid. The left-mastoid was used as reference for all recording sites, and data were re-referenced off-line to the average of all scalp electrodes. The scalp EEG and EOG were amplified with an SA Instrumentation amplifier with a bandpass of 0.01–80 *Hz*, and signals were digitized at 250 *Hz* in LabView 6.1 running on a PC. EEG data was first segmented by 18.5 second intervals to include all trials within a block. After time-frequency decomposition was performed (*see next section*), these epochs were further segmented into trial-to-trial epochs (the time interval of - 600 to 800 ms relative to the onset of the stimulus for both experiments). These trial-to-trial epochs including blinks (>80 μv, window size = 200 ms, window step = 50 ms), large eye movements (>1°, window size = 200 ms, window step = 10 ms), blocking of signals (range = - 0.01 μv to 0.01 μv, window size = 200 ms) were excluded from subsequent analyses.

### Time-Frequency Analysis

Rather than raw EEG, we used the time-frequency decomposed EEG signal for decoding of representations (e.g., Foster, Sutterer, Serences, Vogel, & Awh, 2017; Kikumoto & Mayr, 2018). Temporal-spectral profiles of single-trial EEG data were obtained via complex wavelet analysis (Cohen, 2014) by applying time-frequency analysis to preprocessed EEG data epoched for each block (>18 seconds to exclude the edge artifacts). The power spectrum was convolved with a series of complex Morlet wavelets *e*^2*πft*^ *e*^−*t*2^/(2\**σ*^2^)), where *t* is time, *f* is frequency increased from 1 to 35 *Hz* in 35 logarithmically spaced steps, and σ defines the width of each frequency band, set according to *n*/2*πf*, where *n* increased from 3 to 10. The logarithmic scaling was used to keep the width across frequency band approximately equal, and the incremental number of wavelet cycles was used to balance temporal and frequency precision as a function of frequency of the wavelet. After convolution was performed in the frequency-domain, we took an inverse of the Fourier transform, resulting in complex signals in the time-domain. A frequency band-specific estimate at each time point was defined as the squared magnitude of the convolved signal *Z*(*real*[*z*(*t*)]^2^ + *imag*[*z*(*t*)]^2^) for instantaneous power.

### Representational Similarity Analysis

The decoding analysis in the current study follows closely our previously established methods (Kikumoto & Mayr, 2020). In order to assess the strength of each action feature and conjunction on the level of individual trials and time points, we used a two-step procedure. An initial, linear decoding step yielded similarity information that could be analyzed through the second, representational similarity analysis step. For the initial step, we used a penalized linear discriminant analysis using the caret package in R (Kuhn, 2008) to discriminate between all 12 possible action constellations. At every time sample point, the instantaneous power of rhythmic EEG activity was averaged within the predefined ranges of frequency values (1-3 *Hz* for the delta-band, 4-7 *Hz* for the theta-band, 8-12 *Hz* for the alpha-band, 13-30 *Hz* for the beta-band, 31-35 *Hz* for the gamma-band), generating 100 features (5 frequency-bands X 20 electrodes) to train decoders. Within individuals, these data points were *z*-transformed across electrodes at every time sample to remove the effects that uniformly influenced all electrodes. We used a k-fold repeated, cross-validation procedure to evaluate the decoding results (Mosteller & Tukey, 1968), by randomly partitioning single-trial EEG data into four independent folds. All trials except incorrect go-trials were used as the training sets in both experiments. The number of observations of each action constellation was kept equal within and across folds by dropping excess trials randomly. Three folds served as a training set and the remaining fold was used as a test set; this step was repeated until each fold had served as a test set. Each cross-validation cycle was repeated eight times, in which each step generated a new set of randomized folds. Resulting classification probabilities (i.e., evidence estimated for each case of S-R mapping) were averaged across all cross-validated results with the best-tuned hyperparameter to regularize the coefficients for the linear discriminant analysis. This decoding step yielded for each time point and trial a “confusion-vector” of classification probabilities for both the correct and all possible incorrect classifications (Fig. 1c).

As the second step, we applied time-resolved RSAs to each confusion profile in order to determine the underlying similarity structure. Specifically, we regressed the confusion vector onto model vectors as predictors, which were derived from a set of representational similarity model matrices (Fig. 1c). Each model matrix uniquely represents a potential, underlying representation (e.g., rules, stimuli, responses and conjunctions). For example, the rule model predicts that the decoder would only discriminate instances of different rules, but fail to discriminate instances of the same rule. To estimate the unique variance explained by competing models, we regressed all model vectors simultaneously, resulting in coefficients for each of the four model vectors. These coefficients (i.e., their corresponding *t*-values) allowed us to relate the dynamics of action representations to trial-to-trial variability in behavior during go- and stop-trials (see Multilevel Modeling section for details). For all RSAs, we logit-transformed classification probabilities and further included subject-specific vectors that contained *z*-scored, average RTs and stopping accuracy as nuisance predictors to reduce potential biases in decoding due to idiosyncratic differences in performance among action constellations (see also Fig. S2 and S3 in the Supplemental Material).

We excluded *t*-values that exceeded 5 SDs from means for each sample point, which excluded less than 1% of the entire samples in both experiments. Resulting *t*-values were averaged within 20 ms non-overlapping time samples. For decoding analyses and subsequent RSAs, incorrect go-trials were excluded.

In both experiments, decoders were trained with the stimulus-aligned EEG signal. In Exp. 1, we further computed RSA scores that were re-epoched in reference to the onset of the stop signal (the right column of Fig. 3 and 4). Matching go-trial results were calculated with the SSDs that would have been used if the stop signal appeared in those trials.

### Estimating Timing of Stop-induced Suppression

In Exp. 2, we used nonparametric permutation tests with a single-threshold method to identify the earliest time sample at which statistically significant differences between go-trials and stop-trials emerged. Specifically, for each action feature, we computed permutation distributions of the maximum statistic for every sample point from the stop-signal onset (fixed at 100 ms after the stimulus onset) to the end of 800 ms of the hold period. First, we obtained RSA results by decoding data with randomly shuffled condition labels (i.e., of action constellations). We then performed a series of *t*-tests, testing the differences in RSA scores between go- and stop-trials, for every sample against the null level (i.e., 0 for *t*-values). Out of the series of *t*-test results, we retained the maximum *t*-value. We repeated this process 10000 times by randomly drawing samples from all possible permutations of labels, thereby generating the permutation distributions of the maximum statistics. This approach allowed us to identify statistically significant, individual time points by comparing scores from the correct labels to the critical threshold, which was defined as the 95th (i.e., alpha =.05) of the largest member of maximum statistics in the permutation distribution of the corresponding variable.

### Multilevel Modeling

In Exp. 1, to analyze predictors of trial-by-trial variability in stopping success, we used multilevel logistic regression models. Specifically, we estimated for stop trials a model predicting stopping success on a given trial using the RSA-derived *t*-values for basic action features (i.e., rule, stimulus, and response) and the conjunction as predictors. In addition, we also included each trial’s log-transformed SSD as a covariate to account for the possibility that SSDs affect both action representations and stopping success as a third-variable. For statistical tests, we used EEG data averaged over a-priori selected, symmetric time intervals, namely a pre-stop-signal (−200 to 0 ms) and a post-stop-signal period (0 ms to 200 ms), relative to the onset of the stop signal in each trial. Both time intervals clearly precede the average SSRT across individuals (M = 272 ms). We also performed additional control analyses, where we excluded trials with early responses (i.e., responses occurred after the stimulus onset and before the stop signal in unsuccessful-stop trials) and where we included decoded representations from both pre- and post-stop-signal phases simultaneously (Table 3). In addition, to visualize changes in predictability of stopping success, we separately performed a series of logistic regression analyses by fitting models at each sample point in reference to the onset of the stimulus and the stop signal (Fig. 5). In order to replicate the results by (Kikumoto & Mayr, 2020) about how action representations contributed to action selection in go-trials, we also report for both experiments results from multilevel models to assess which action representations predict trial-to-trial RTs on go trials (Table 2). Here, RTs were log-transformed and trials with response errors were excluded.

### Open Practice Statement

Neither of the experiments reported in this article was formally preregistered. All data and analysis scripts will be posted on OSF for the final manuscript.

## Results

### Experiment 1

#### Behavior

Behavioral performance is summarized in Table 1. Most participants (33 out of 36 participants) exhibited *p*(stop|signal) in the range of .40-.65; individuals with the stopping accuracy higher than 75% were excluded from further analyses. The average RTs in go-trials were longer than the RT in failed stop-trials for all participants. This pattern is consistent with the race model as a basis for estimating individuals’ stop-signal reaction time (SSRT, Fig. 2a). Also, the probability of stopping errors covaried with the increase of SSDs, indicating the overall efficacy of the SSD staircase algorithm (Fig. 2b).

**Fig. 2.**
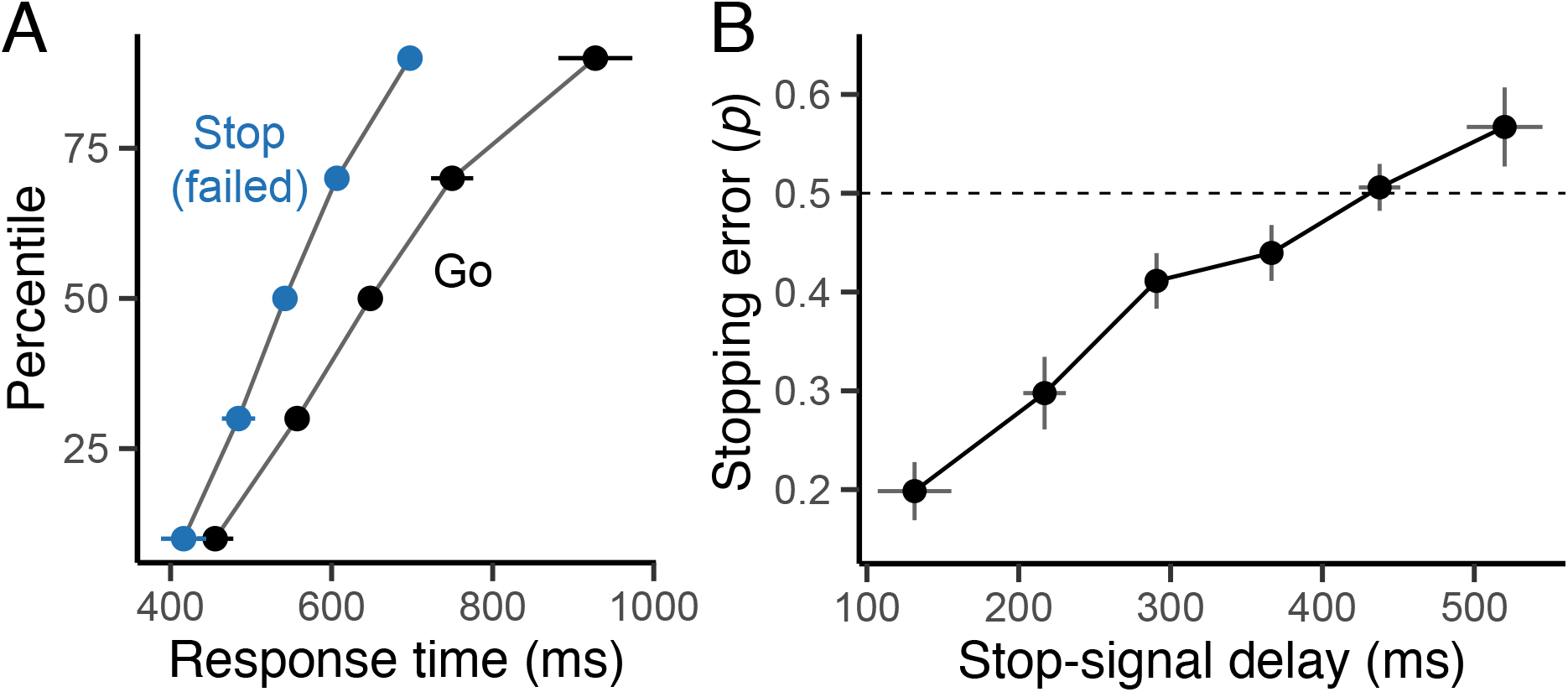
Behavioral results in Exp. 1. (A) Vincentized mean response times (RTs) for go-trials and failed stop-trials. (B) Average rates of stopping failures as a function of stop-signal delays. Error bars specify 95% within-subject confidence intervals.

**Table 1.**
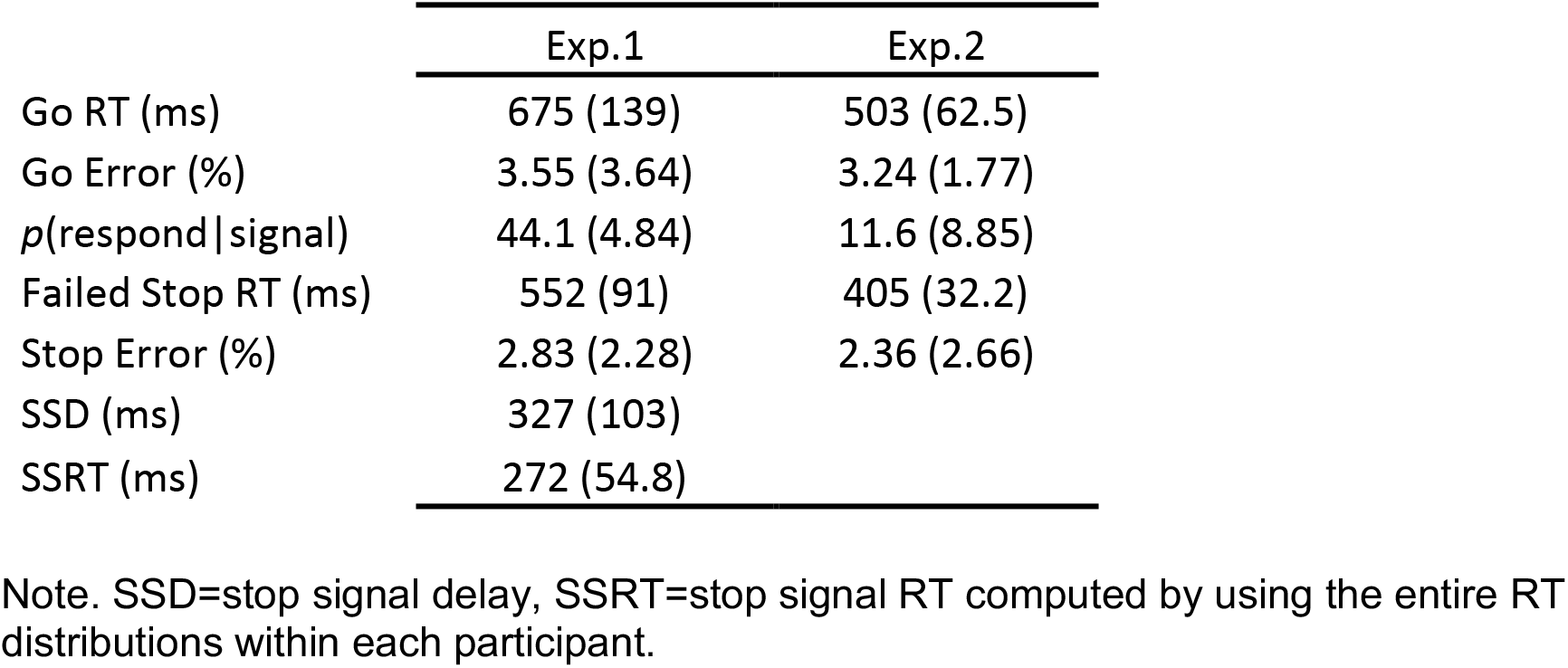
Behavioral performance in go- and stop-trials.

#### Action Representations in Go-trials, Failed Stop-trials, and Successful Stop-trials

Fig. 3 shows the time-course of RSA scores estimated on the level of single trials for each of the basic features (i.e., rules, stimuli, and responses) and the conjunction all trial types. For go-trials, the flow of activated representations was highly consistent with our previous results. Rule information appeared in the pre-stimulus period, stimulus information peaked shortly after the stimulus appeared, followed by the emergence of response information (Hubbard, Kikumoto, & Mayr, 2019; Kikumoto & Mayr, 2020). Importantly, conjunctive information was present throughout the entire response-selection period. We also replicated the previous finding that trial-to-trial variability in conjunctive representations robustly predicted go-trial RTs (Table 2), over and above other representations of constituent features. Note, that in the Supplemental Material, we include an additional analysis that probes the robustness of the decoding/RSA analyses.

**Fig. 3.**
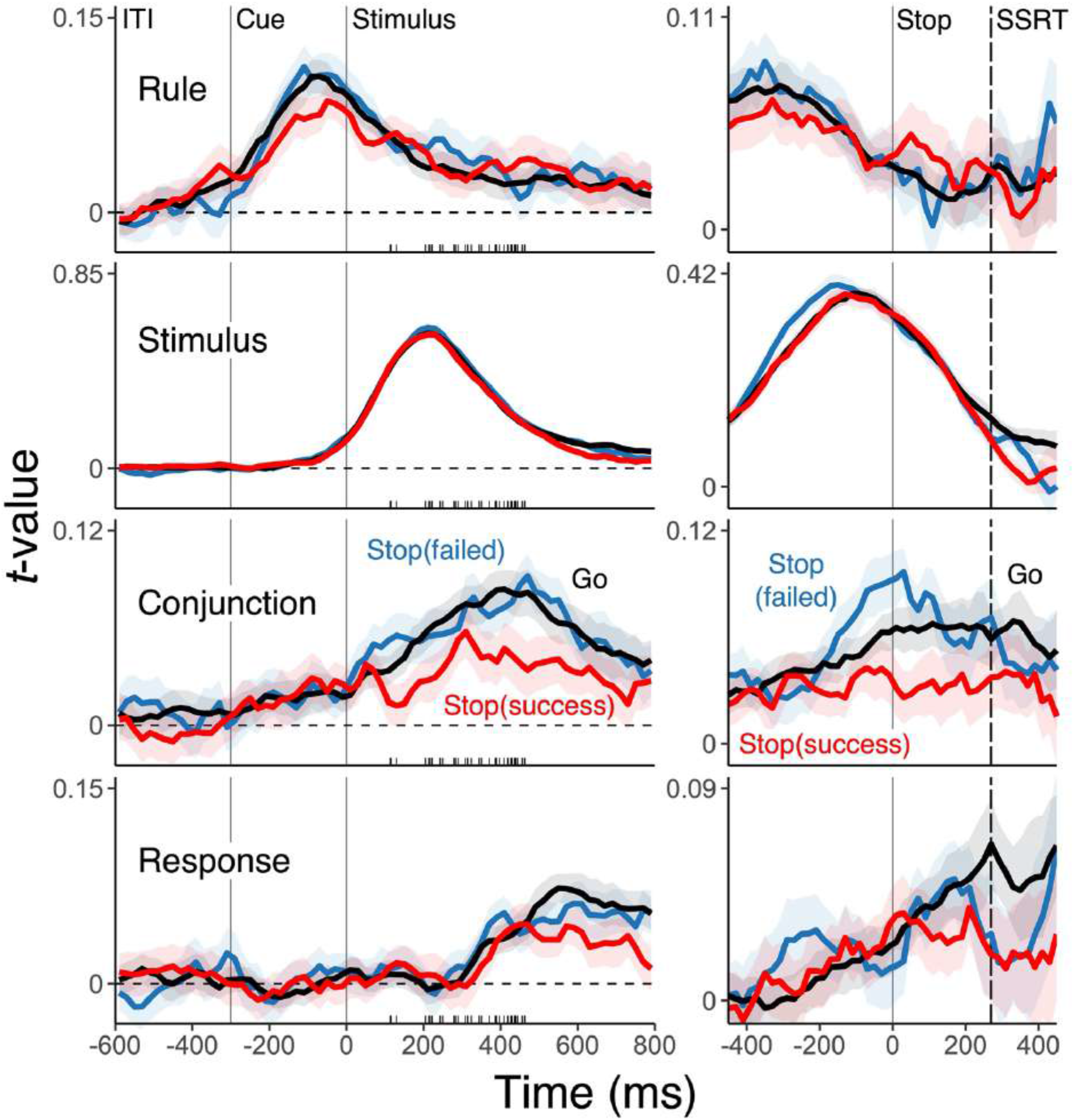
Effects of stopping on representations in successful- and failed-stopping trials compared to go-trials. Average, single-trial *t*-values associated with each of the basic features (rule, stimulus, and response) and their conjunction derived from the RSA, separately for go-trials (black), successful stop-trials (red), and failed stop-trials (blue). The left panels show the results aligned with the stimulus onset, the right panels aligned with the stop-signal onset. Shaded regions specify the 95% within-subject confidence intervals. Tick marks on the x-axis of the stimulus-aligned panels mark individuals’ average stop-signal delays.

**Table 2.** Predicting trial-by-trial RTs in go-trials using the average strength of representations decoded through the RSA analyses during 0-300 ms post-stimulus intervals for each trial.

| Variable | Exp.1 |  | Exp.2 |  |
| --- | --- | --- | --- | --- |
|  | *b* (se) | *t*-value | *b* (se) | *t*-value |
| Rule | -.013 (.006) | -2.08 | -.025 (.012) | -2.15 |
| Stimulus | -.038 (.009) | -4.12 | -.015 (.009) | -1.59 |
| Response | -.032 (.009) | -3.55 | -.016 (.009) | -1.85 |
| Conjunction | -.057 (.010) | -5.69 | -.042 (.012) | -3.61 |

Note. Coefficients for all decoded variables were included as predictors simultaneously. Negative coefficient imply that stronger representations predict faster RTs.

The strength of conjunctive representations and late response representations were selectively reduced in successful stop-trials relative to go-trials and failed stop-trials. In contrast, there were no clear effects on rule and stimulus representations. For the conjunctive representations, the divergence between failed and successful stop trials occurred even before the onset of the stop signal (see individuals’ average SSDs in Fig. 3 left column), suggesting stopping was particularly impaired when the conjunctive representations were strong in the early response selection phase. Indeed, when we replotted RSA scores relative to trial-to-trial SSDs (see Fig. 3, right column), differences in successful and failed stop-trials emerged clearly before the average SSRT (M = 272 ms) and even before the stop signal. No other action features showed similar differences in the pre-stop-signal period (*t* < .13), and post-stop-signal effects on the response representation were apparent only when the conjunction model was excluded, *b* = −.010, SE = .004, *t*(33) = 2.34.

Before accepting the conclusion that the state of conjunctive representations prior to the onset of the stop signal determined the success of stopping, we need to consider the fact that trial-to-trial variability in SSDs is likely to covary with both the strength of conjunctions and the probability of successful stopping. Therefore, to rule out SSDs as a potential third-variable explanation, we performed multilevel logistic regressions to predict single-trial stopping failures using decoded action features and SSDs as simultaneous predictors. Fig. 4, shows time-point by time-point results of these analyses, which clearly indicate that pre-stop-signal conjunctions are a unique predictor of stopping success. Statistical tests of these relationships confirmed that the average state of conjunctions prior to the onset of the stop signal strongly predicted stopping failures over and above the state of the other feature representations (Table 3, top panel). To ensure that these results are not due to very fast responses that occurred prior to the stop signal, we confirmed that these results were robust when eliminating premature responses (Table 3, middle panel). In addition, when entering pre-stop-signal (−200 ms – 0 ms) and post-stop-signal predictors (0 ms – 200 ms) simultaneously, we found that during each phase, conjunctions uniquely predicted stopping success (Table 3, bottom panel).

**Fig. 4.**
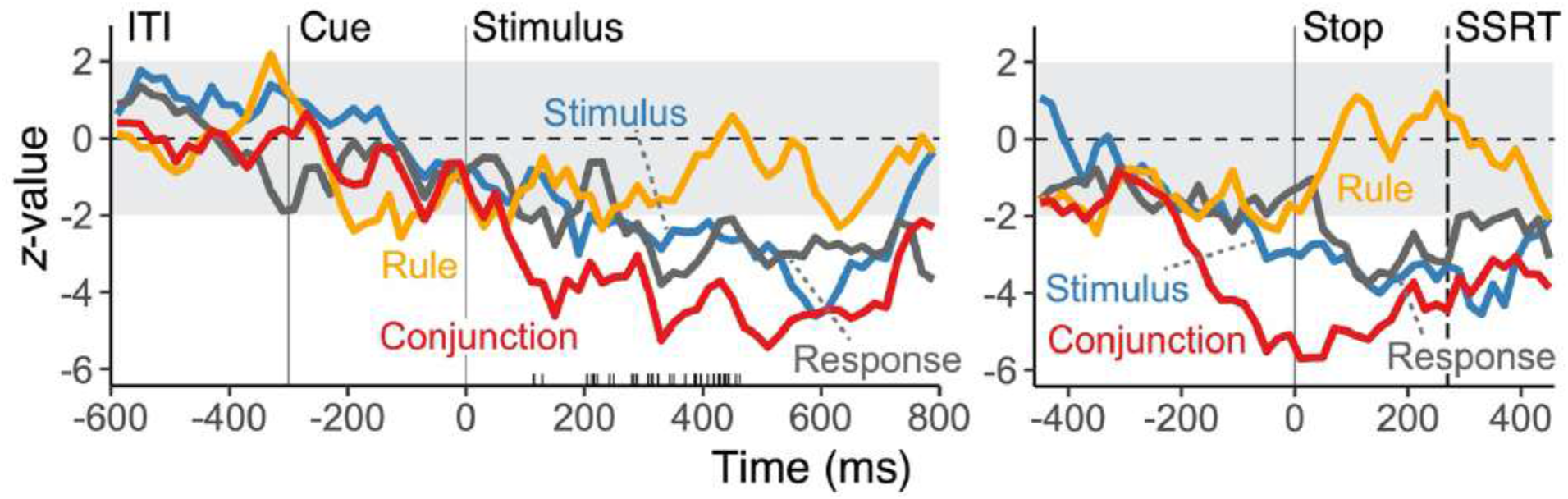
Predicting stopping success from strength of representations. Time-course of *z* values from multilevel, logistic regression models predicting the variability in trial-to-trial stopping failures in the stop-trials (the “impact” of representations on stopping success), using RSA scores of all features and trial-to-trial SSDs as simultaneous predictors. Negative *z*-value indicates more stopping failures as the strength of decoded representations increase. The left panel shows results aligned to the stimulus onset; in the right panel data are aligned to the stop-signal onset. Tick marks on the x-axis of the stimulus-aligned panels mark individuals’ average stop-signal delays.

**Table 3.** Predicting trial-by-trial stopping accuracy using the strength of decoded representations in Exp. 1.

| Model | Variable | Pre-Stop-Signal |  | Post-Stop-Signal |  |
| --- | --- | --- | --- | --- | --- |
|  |  | *b* (*se*) | *t*-value | *b* (*se*) | *t*-value |
| SSD control | Rule | -.077 (.033) | -2.38 | -.012 (.024) | -0.49 |
|  | Stimulus | -.083 (.033) | -2.50 | -.060 (.020) | -3.11 |
|  | Response | -.081 (.034) | -2.41 | -.062 (.020) | -3.11 |
|  | Conjunction | -.193 (.041) | -4.70 | -.151 (.029) | -5.23 |
| Exclude early responses | Rule | -.050 (.020) | -2.43 | -.019 (.025) | -0.76 |
|  | Stimulus | -.036 (.017) | -2.19 | -.057 (.020) | -2.89 |
|  | Response | -.039 (.021) | -1.86 | -.054 (.020) | -2.70 |
|  | Conjunction | -.136 (.029) | -4.78 | -.155 (.029) | -5.31 |
| Pre/post stop signal simultaneous | Rule | -.084 (.036) | -2.31 | .016 (.044) | 0.36 |
|  | Stimulus | -.035 (.036) | -0.93 | -.104 (.041) | -2.53 |
|  | Response | -.038 (.040) | -0.97 | -.093 (.041) | -2.30 |
|  | Conjunction | -.126 (.049) | -2.60 | -.194 (.049) | -3.93 |

Note. The “SSD control” model included trial-to-trial stop signal delays as the fixed and random effect as covariate. In the “exclude early responses” model, all premature responses that occurred prior to the onset of the stop signal were removed. Whereas these models were fitted separately for pre-stop-signal and post-stop-signal predictors, in the “pre/post stop signal simultaneous” model, both pre-stop-signal and post-stop-signal predictors were included simultaneously. Pre-stop-signal interval: -200–0 ms; post-stop-signal interval: 0-200 ms.

Arguably, a pre-stop-signal effect of the conjunctive representations may be driven by participants who used occasionally used a strategy of waiting for the stop signal in order to initiate the go process. We used a median split to separate subjects by the proportion of stopping failures. Both pre- and post-stop-signal states of conjunctions predicted stopping failures even in the below-median group, who should be least likely to use a waiting strategy (Table S1). This indicates that the observed results cannot be attributed to a subset of particularly cautious participants.

The stop-signal-aligned pattern shown in Fig. 3 (right column) is also suggestive of conjunctions as a representational target of the stopping process. Specifically, in failed-stop trials, conjunctive representations were particularly strong at the time the stop signal arrived, but immediately following the stop signal showed a rapid reduction. Such a pattern is consistent with a stop-signal-induced suppression, but that came too late to influence behavior. When analyzing the 320 ms period following the stop signal (using eight 40 ms bins), the linear trend was indeed stronger for failed-stop compared to successful-stop or go trials combined, b = .008, SE = .0038, *t* = 2.04. However, this pattern is also somewhat inconclusive as the reduction following the peak at the time of the stop signal might also be interpreted as a regression towards the mean level of conjunction strength. In Exp. 2, we will seek more definitive evidence regarding the representational targets of the stopping process.

### Experiment 2

Exp. 1 clearly demonstrated that the strength of conjunctive representations is strongly predictive of trial-to-trial stopping success and also presented initial evidence that conjunctive representations are predominantly affected by the stopping process. However, while the adaptive calibration of the stopping success rate around 50%, was useful for determining the relationship between conjunctive representations and behavioral performance, it made it more difficult to distinguish between the representational precursors and consequences of stopping success. In order to clearly establish the effects of stopping on different representations, we used a consistent, early stop signal in Exp. 2, which ensured high, overall stopping success.

#### Behavior

Behavioral performance in go/stop-trials are summarized in Table 1. The probability of stopping failures—incorrectly executing responses in the presence of the stop signal (i.e., *p*(respond|signal))—was low because of the early presentation of stop signal at the fixed timing (100 ms after the stimulus onset). This allowed us to estimate the time-course of suppression of action representations from a fixed starting point. Note, that the substantially lower RTs for go trials in Exp. 2 compared to Exp. 1, are likely due to the reduced uncertainty about the timing of the stop signal in Exp. 2.

#### Action Representations in Go-trials and Stop-trials

As shown in Fig. 5, the pattern of activated representations was highly consistent with our previous results. Also again, conjunctive representations robustly predicted go-trial RTs (Table 2), over and above representations of constituent features.

**Fig. 5.**
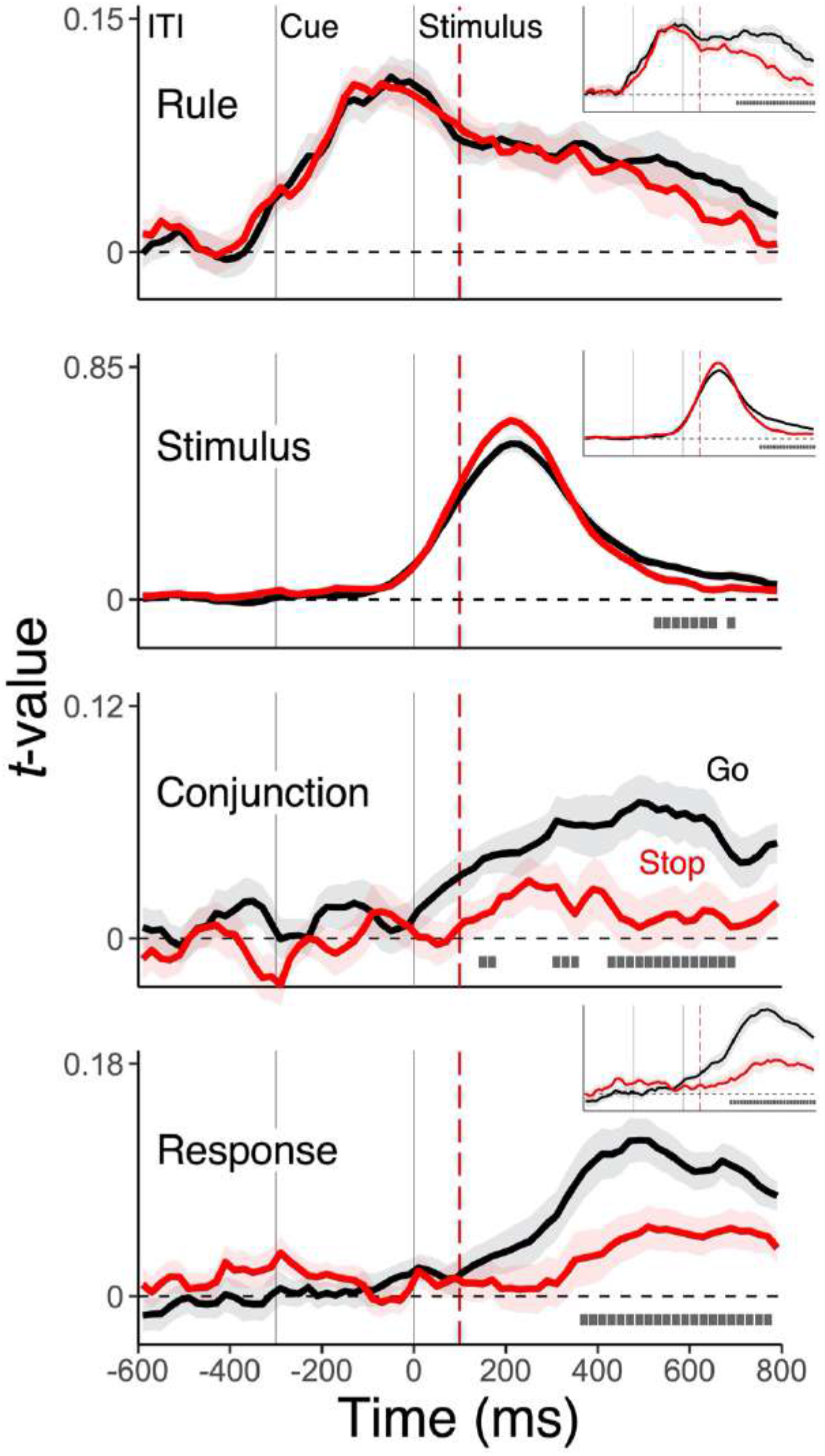
Effects of stopping on representations in the fixed-stop delay. Average, single-trial *t*-values derived from the RSA (see Fig. 1C) for each of the basic features (rule, stimulus, and response) and the conjunction, separately for go-trials (black) and stop-trials (red). Shaded regions specify the 95% within-subject confidence intervals. The vertical, red dashed line marks the onset of the stop signal at 100 ms after the stimulus onset. Gray squares below lines denote the time points with significant differences between go- and stop-trials, correcting for multiple comparison using a non-parametric permutation test. The inserts for the rule, stimulus, and response features show the same results when the RSA contains only these basic features, but excludes the conjunction as model predictor.

Our main goal in Exp. 2 was to test the prediction that conjunctive representations are suppressed on stop-trials relative to go-trials. Indeed, we found stopping of actions markedly reduced the strength of conjunctive representations right after the onset of the stop signal (Fig. 5). Not surprisingly, the response representation was also suppressed, whereas we found no effect on the rule representation and only a late, and small effect for the stimulus representation. Yet, when only the basic constituent features (i.e., rules, stimuli and responses) were used in the RSA model, the suppression effect was substantially increased for the rule representation, highlighting the importance of including the conjunction model (see also, Kikumoto & Mayr, 2020). Suppression of the conjunction occurred at the same time, or even slightly before suppression of the response representation. This suggests that the reduction of conjunctive information is not just an aftereffect of response suppression. Rather, it supports the notion that the conjunctive representation is a direct target of the stopping activity.

## Discussion

Even simple, goal-directed actions rely on various aspects of the task environment. Both psychological and neural-level theories of action propose that all relevant action features need to be integrated within conjunctive representations for successful action selection (Hommel et al., 2001; Rigotti et al., 2013). Using a convenience sample of student participants, we tested here the hypothesis that because such representations are critical for action selection, they should also be intricately involved when a planned or initiated action needs to be stopped. Consistent with this hypothesis we found that the strength of conjunctive representations at the time the stopping process is initiated, inversely predicts stopping success (Fig. 3 and 4) and that conjunctive representations are a main target of the stopping process (Fig. 3 and 5).

In principle, stopping of actions might require suppression of all task-relevant representations. Alternatively, only those representations directly involved with motor control might be targeted. Instead, our results are most consistent with the hypothesis that the conjunctive representation is the primary target of suppression, followed by the response representation (Fig. 3, 4, and 5). It is an open question whether conjunctive and response representations are separately targeted, or whether the deactivation of response representations is a consequence of the suppressed conjunctive representations. Representations of the rule or the stimulus remained intact, or showed very minor suppression, and only after the completion of the stopping process (Fig. 3 and 5). A potential functional benefit of selective suppression in real-world situations is that by preserving the rule information, actions can be easily re-implemented, once the reason for stopping has been removed.

Conjunctive representations that integrate stimulus, response, and rule information are by definition situated on a more central level than representations that directly control motor output. The fact that conjunctive representations were targeted by the stopping process, is consistent with results indicating that inhibition of actions and inhibition of thoughts or memories are handled by a shared process (Anderson, 2004; Guo, Schmitz, Mur, Ferreira, & Anderson, 2018). For example, using the Think/No-think paradigm, studies found that the same right lateral prefrontal area that is typically involved in stopping of motor responses, was also critical in suppressing thoughts, leading to longer-term negative effects on their accessibility. One might speculate that conjunctive representations as a target of inhibition may mitigate such longer-term effects of suppression (Anderson & Green, 2001). Specifically, by adding contextual specificity to abstract feature codes, conjunctive representations should help constrain inhibition to the currently relevant context.

Our results showed that the pre-stop-signal state of the conjunctive representation uniquely predicts the success of subsequent stopping, over and above the potential effects of a reactive inhibition process that is initiated after the stop signal (Table 3). Thus, the strength of conjunctive representations is a key driver of the efficiency and success of an action—and therefore also of the ability to stop that action. However, our results by themselves do not identify the underlying mechanisms that modulate the state of the conjunctive representation prior to the stop signal. One possibility is that conjunction strength depends on endogenous fluctuations of attention towards the go-action across trials. A strong emphasis on initiating action may induce strong conjunctions and thereby cause the failures to trigger the stop process altogether (Matzke, Hughes, Badcock, Michie, & Heathcote, 2017; Matzke, Love, & Heathcote, 2017). The fact that conjunctive representations in failed-stop trials, prior to the arrival of stop signal, were even stronger than on go-trials, is consistent with such an attentional fluctuation account. As another, not necessary mutually exclusive possibility, there is evidence that variations in strategic, proactive inhibition (Aron, 2011) that may affect the state of conjunctions. On trials in which subjects anticipate stopping, proactive inhibition may keep conjunctive representations from fully developing. Such proactive control processes could be directly tested, by cuing the stop probability on a trial-by-trial basis (Chikazoe et al., 2009; Vink, Kaldewaij, Zandbelt, Pas, & du Plessis, 2015; Zandbelt, Bloemendaal, Neggers, Kahn, & Vink, 2013). In any case, our results clearly confirm that the pre-stop-signal state of action representations must be taken into account to fully understand subsequent reactive inhibition and stopping.

One potential caveat is that for the task space used in Exp. 1 and 2, the RSA analyses allow us to say with certainty that at least two different task-relevant features were integrated within conjunctions, but do not allow disambiguating between conjunctions that include binary combinations of rules, stimuli, or responses, or the combination between all three aspects. However, previous work had also used an expanded task space that allowed disambiguating between different types of conjunctions (Exp.2 in Kikumoto & Mayr, 2020). In terms of functional characteristics, the rule-independent (i.e., stimulus-response) conjunctions from the limited task space, and the rule-specific conjunctions from the extended task space, had behaved in a highly similar manner. In addition, our current results showed that when the conjunction predictor was dropped from the RSA analyses, the coefficients for the rule predictor absorbed much of the conjunction effect—indicating a contribution of rule information to the decoded conjunctions (Fig. 5). Therefore, the observed conjunctions likely reflect an integration between both the rule, and stimulus/response features.

Our EEG-based decoding results provide no precise information about the neural-anatomical location of conjunctive representations (for related exploratory analyses, see supplementary information to Kikumoto & Mayr, 2020). However, recently there has been increasing evidence from research with non-human primates about the high prevalence of neurons in parietal/frontal areas that show very similar properties as the conjunctive representations we report on (Fusi, Miller, & Rigotti, 2016; Parthasarathy et al., 2017; Rigotti et al., 2013). Specifically, these so-called mixed-selectivity neurons integrate various task-features in a nonlinear and diverse manner and, just as the EEG-decoded conjunctive representations, are uniquely predictive of successful action selection. It would be important to establish to what degree the conjunctive representations examined here, reflect mixed-selectivity, neuronal activity. One way to test this hypothesis is to look for equivalent functional and computational properties of both conjunctive and mixed-selectivity representations in both human and animal models (Badre, Bhandari, Keglovits, & Kikumoto, 2020; Bernardi et al., 2020).

In conclusion, the results we report here build on our previous work suggesting that conjunctive, event-file type representations can be tracked and related to behavior with high temporal resolution through EEG-decoding techniques. Specifically, our results are consistent with the hypothesis that such conjunctive representations are a prime target of action inhibition, exactly because they are a key driver of successful action implementation.

## Acknowledgements

This research was supported by NIA grant R01 AG037564-01A1, and by NSF grant NSF grant 1734264.

## SUPPLEMENTAL MATERIAL

### Event-related potential (ERP) to the stop signal in Exp. 1

A number of previous studies have analyzed ERPs in the context of versions of the stopsignal paradigm. Typically, a frontocentral N2/P3 complex has been found to be affected in response to the stop signal (Greenhouse & Wessel, 2013; Kok, Ramautar, De Ruiter, Band, & Ridderinkhof, 2004; Wessel & Aron, 2015). There is some debate about the cognitive processes reflected by these different ERP measurements. Nevertheless, we wanted to confirm that we can replicate the standard ERP pattern in our paradigm, which is considerably more complex than standard stop-signal tasks. Consistent with Wessel and Aron (2015), we observed a pattern of ERPs, where the onset latency of the frontocentral P3 was shorter in successful stop-trials compared to failed stop-trials (Fig. S1).

**Fig. S1.**
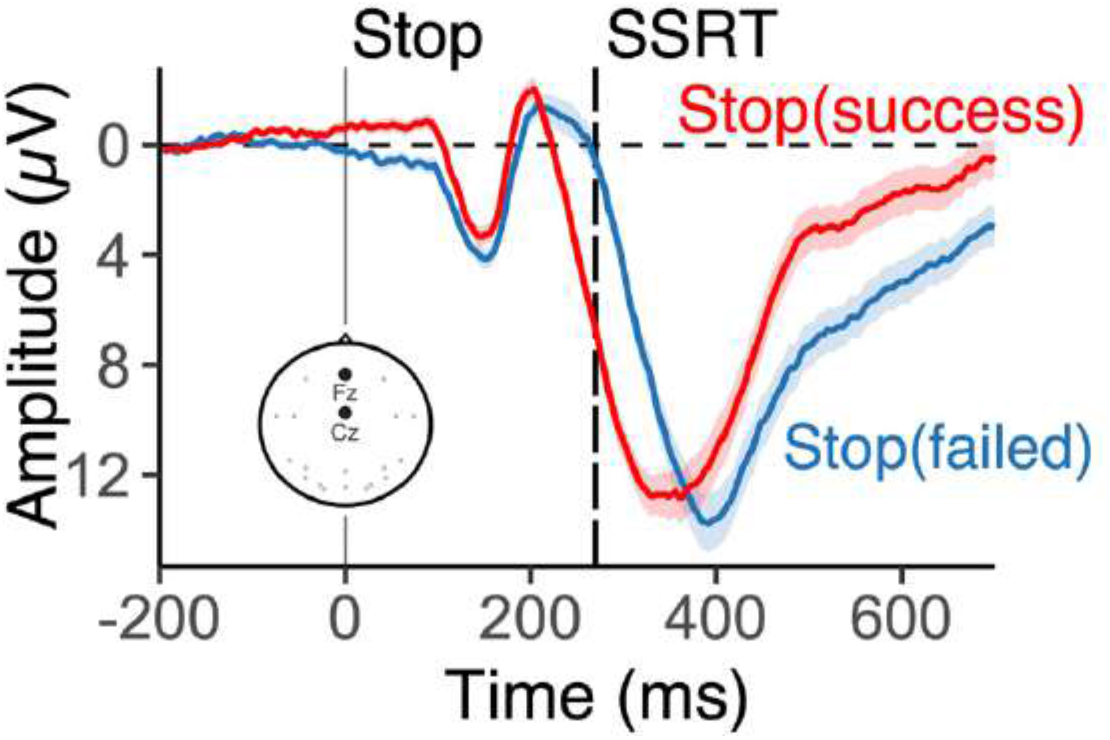
Grand averaged ERPs for successful and failed stop trials, time-locked to the onset of trial-to-trial stop signal.

### Predicting single-trial stopping failures

We observed that both pre- and post-stop-signal states of conjunctive representations predict stopping failures at the level of single trials. Yet, one potential concern is that at least some participants may have engaged in a waiting strategy and that this group of subjects in particular is responsible for the predictive relationship between pre-stop-signal conjunction strength and decoding accuracy. Therefore, we repeated our main multi-level logistic regression analyses after separating participants into a low stopping failure group (i.e., potentially using a waiting strategy) and high stopping error group. Importantly, both groups showed the predictive relationship linking conjunctions to stopping success.

**Table S1.**
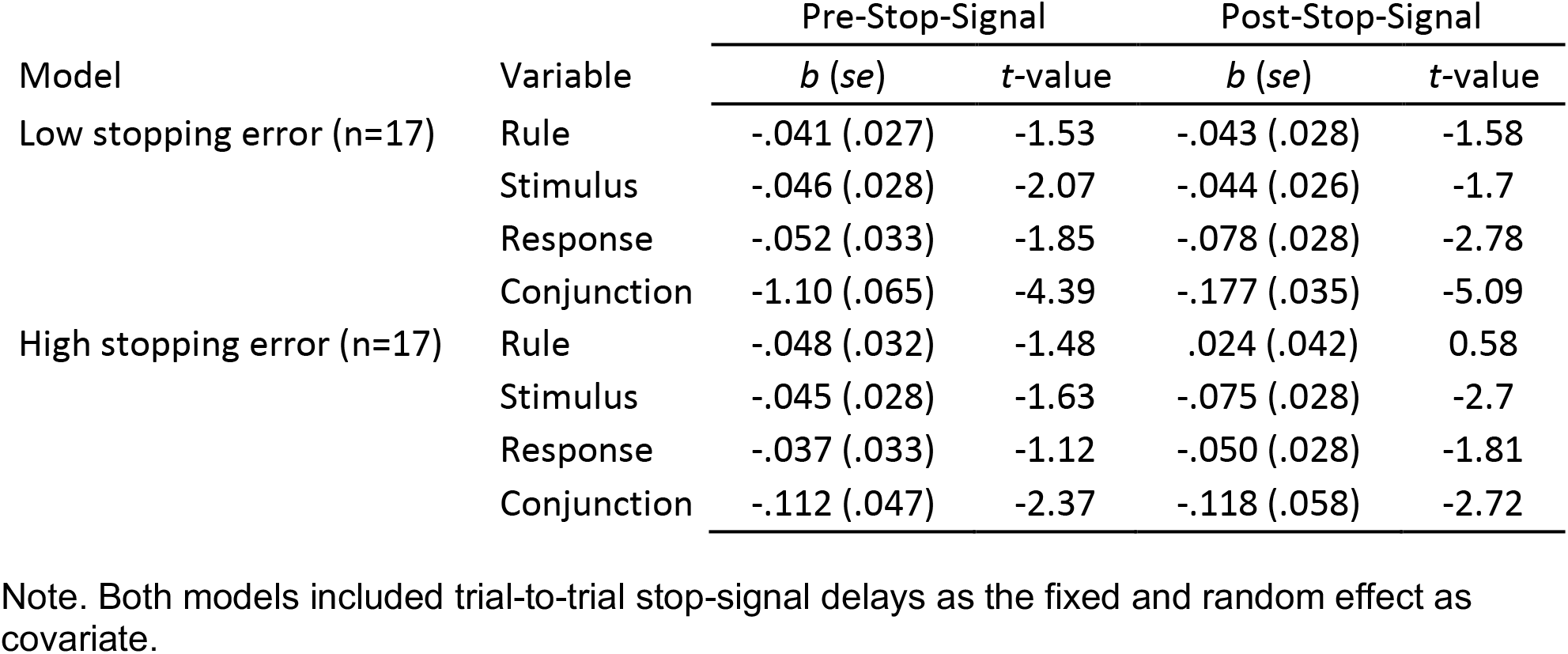
Predicting trial-by-trial stopping accuracy using the strength of decoded representations in Exp. 1.

### Individual variability in stopping performance

Fig. S2 and S3 show the average and individual-specific variability in RT and stopping errors across all rule-stimulus-response configurations for both Exp. 1 and 2. These individual-specific profiles of two behavioral measures were added as additional predictors into the RSA analyses in order to control for difficulty effects specific to each constellation. RSA results without these control predictors were similar to the ones reported in the main text.

**Fig. S2.**
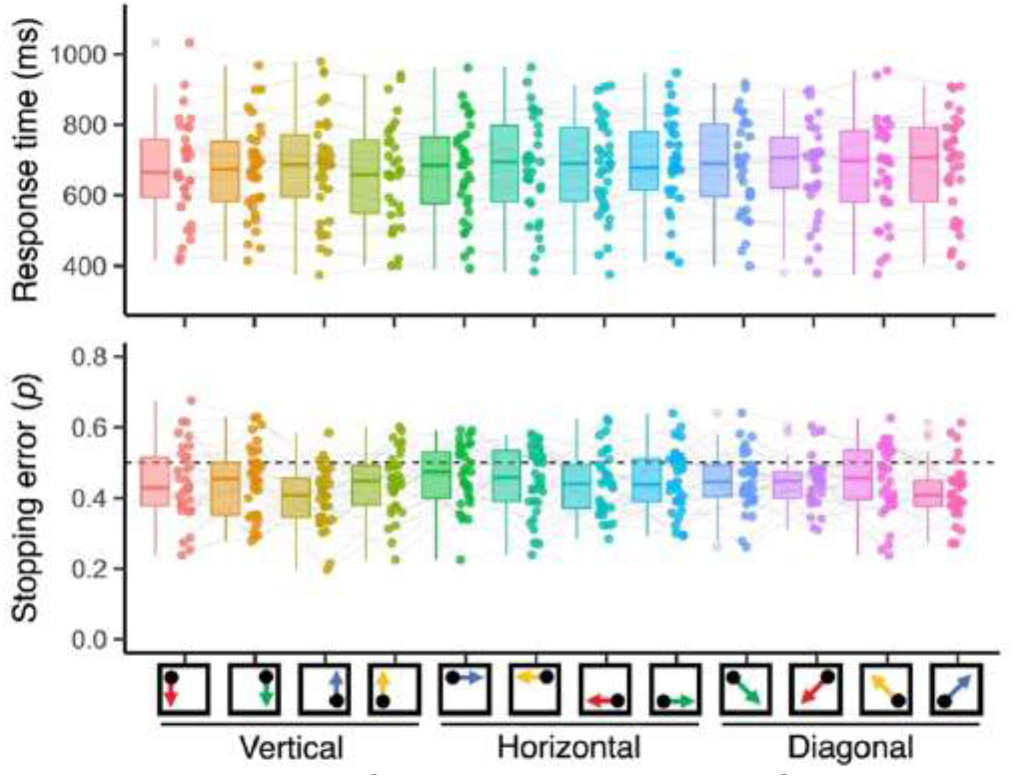
Mean RTs and stopping errors of individual subjects for all action constellations in three different rules. Using a adaptive tracking method, stopping errors converged to .5 on average. To control potential differences covarying with average RTs and stopping errors, vectors of subject-specific RTs and stopping errors were included as nuisance predictors during RSA fitting.

**Fig. S3.**
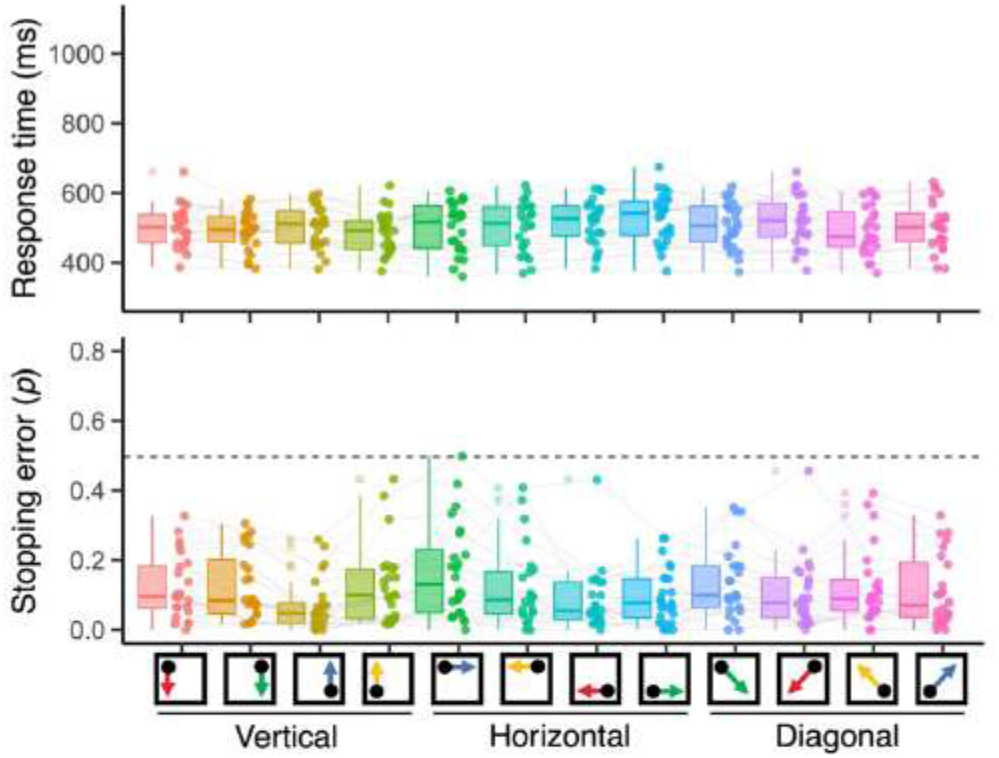
Mean RTs and stopping errors of individual subjects for all action constellations in three different rules. With a fixed stop-signal delay (100ms after the onset of stimulus), stooping errors are reduced on average. To control potential differences covarying with average RTs and stopping errors, vectors of subject-specific RTs and stopping errors were included as nuisance predictors during RSA fitting.

### Evaluating the potential bias in single-trial RSA of conjunction

Our approach to decode the conjunctive representation via single-trial RSA assumes that the decoding results reflect the similarity among action contexts in an unbiased manner. Because decoders are optimized to discriminate each of the possible action constellations from each other, it is possible that superfluous factors affecting discriminability might inflate coefficients of the conjunction regressor (e.g., overfitting of noise). Our theoretically critical results rely on comparisons between conditions (e.g., between stop and go trials), not on the absolute level of coefficients. Nevertheless, we present here a control analysis to examine the potential role of biases introduced during the decoding step.

One way to eliminate or reduce potential bias is to train decoders to classify feature constellations within an “expanded” task space that includes as a subset the conjunctions defined by the combination of rules and stimuli/responses. Specifically, we trained decoders to discriminate 24 instances of action constellations that were defined by the combination of 3 rules, 4 S-R links, and even/odd experimental blocks. Here the decoder would be optimized towards discriminating the 24 different rule/S-R/odd-even conjunctions, and not between the theoretically critical rule/S-R conjunctions. If a decoding bias has a major effect on results, we would expect that the coefficients for the rule/S-R conjunctions are reduced compared to the analysis using the original task space with 12 rule/S-R constellations (see Fig. 4 and 5). However, we found that single-trial RSAs in the expanded space produced almost identical results in both Exp. 1 and Exp. 2 (Fig. S4) to the ones reported in the main text (Fig. 4 and 5). Thus, it is unlikely that the original evidence regarding conjunctive representations is affected by a bias introduced during the decoding step.

**Fig. S4.**
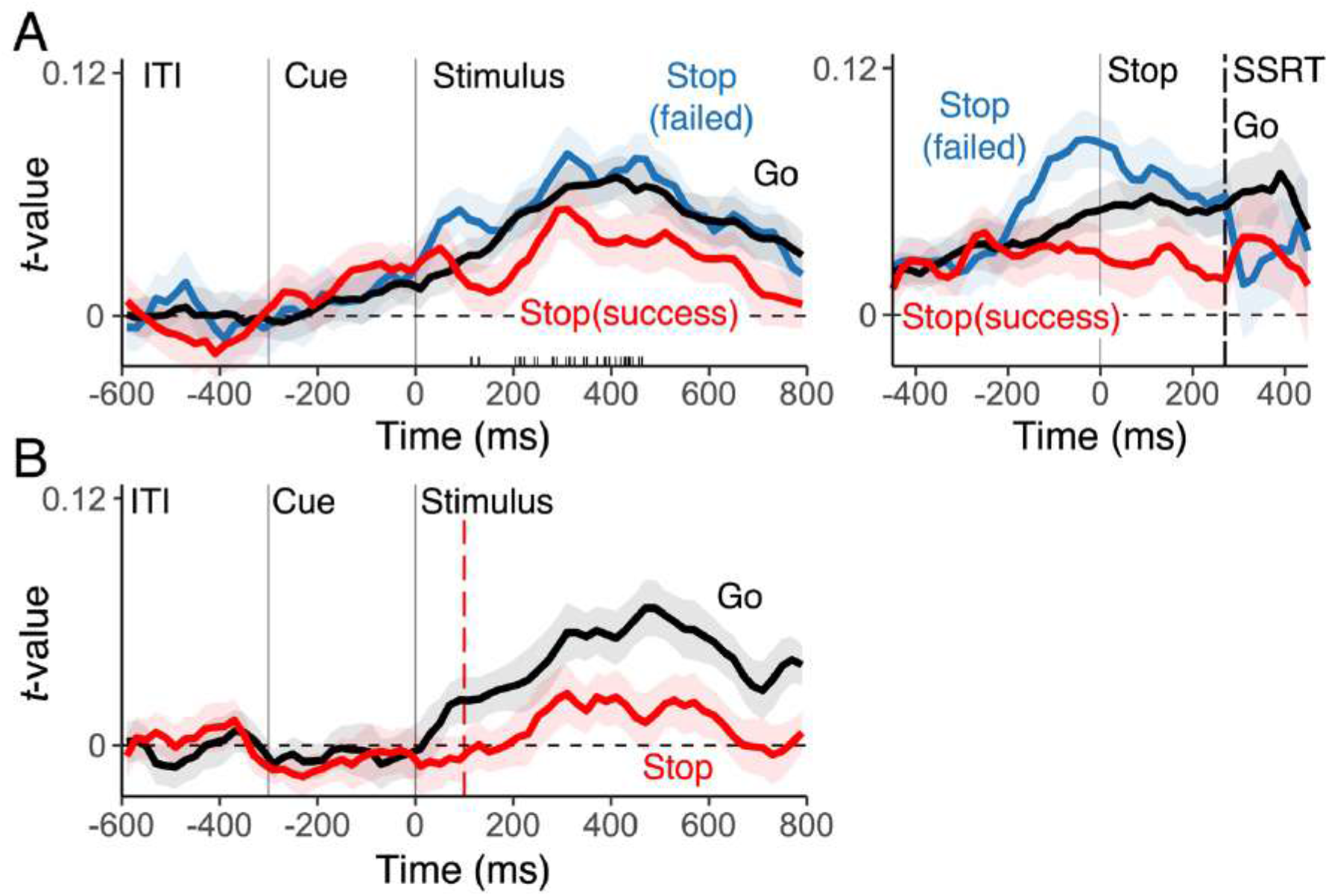
(A) Using the expanded space method, the average, single-trial *t*-values of the conjunction in Exp.1 separately for go-trials (black), successful stop-trials (red), and failed stop-trials (blue). The left panels show the results aligned with the stimulus onset, the right panels aligned with the stop-signal onset. (B) The conjunctive representation in Exp. 2. We present here only the results for conjunction representations; results for the remaining features are also virtually indistinguishable from the results presented in the main paper.

## Notes

### Competing Interest Statement

The authors have declared no competing interest.

### Summary of Updates

Added control analyses and SI.

